# The longevity response to warm temperature is neurally controlled via the regulation of collagen genes

**DOI:** 10.1101/2021.09.26.461885

**Authors:** Sankara Naynar Palani, Durai Sellegounder, Yiyong Liu

## Abstract

Studies in diverse species have associated higher temperatures with shorter lifespan and lower temperatures with longer lifespan. However, the mechanisms behind these inverse effects of temperature on longevity are not well understood. Here, we demonstrate that in *Caenorhabditis elegans*, functional loss of NPR-8, a G protein-coupled receptor related to mammalian neuropeptide Y receptors, increases worm lifespan at 25°C but not at 20°C or 15°C, and that the lifespan increase at 25°C is regulated by the NPR-8-expressing AWB and AWC chemosensory neurons as well as AFD thermosensory neurons. RNA sequencing revealed that both warm temperature and old age profoundly alter gene expression. Further investigation uncovered that the NPR-8-dependent longevity response to warm temperature is achieved by regulating the expression of a subset of collagen genes. As elevated collagen expression is a common feature of many lifespan-extending interventions and enhanced stress resistance, collagen expression could be critical for healthy aging.

## Introduction

Studies in diverse species have associated lower temperatures with longer lifespan and higher temperatures with shorter lifespan, aside from extreme temperatures that could endanger organism survival (*1*). These inverse effects of temperature on longevity were first recorded in poikilotherms, such as the fruit fly *Drosophila melanogaster* (*2*), the cladoceran crustacean *Daphnia magna* (*3*), the nematode *Caenorhabditis elegans* (*4*), as well as in poikilothermic vertebrates such as the South American fish *Cynolebias* (*5*). This phenomenon was also observed in homeotherms such as mice (*6*) and humans (*7*), although with some obscurity (e.g., females have higher body temperatures than males and yet females live longer (*1*)). Nonetheless, there is a clear trend for higher temperatures being associated with shorter lifespan across species.

Most early studies on the longevity response to temperature attempted to interpret the inverse relationship using the rate of living theory, which posits that higher temperatures increase chemical reaction rates, thus speeding up the aging process, whereas lower temperatures do just the opposite (*8*). At the core of this theory is applying the second law of thermodynamics to living organisms, speculating that aging results from increased molecular disorder, and that higher temperatures enhance thermodynamic entropy, thus inducing a faster rate of aging (*8*). Although temperatures could alter the accumulation of molecular damage, recent studies have identified specific cells, cellular molecules, and signaling pathways that are involved in the longevity response to temperature, indicating that this response is a regulated process and not simply due to temperature-induced changes in thermodynamics (*1, 9*). In *C. elegans*, the cold-sensitive TRP channel TRPA-1 can detect temperature drops in the environment and extend lifespan by inducing calcium influx into the cell and activating protein kinase C, which, in turn, activates the pro-longevity transcription factor DAF-16/FOXO (*10*). Interestingly, TRPA-1 also mediates lifespan shortening when worms are subjected to low-temperature treatment during the larval stage (*11*). Moreover, thermosensory AFD neurons and AIY interneurons were found to be required for maintaining *C. elegans* lifespan at warm temperature (25°C) through the DAF-9/DAF-12 steroid-signaling pathway (*12*). Such neural regulation was controlled by the worm cyclic AMP-responsive element-binding protein (CREB) transcription factor (*13*). DAF-41, the *C. elegans* homolog of co-chaperone P23, which has a possible role in protein folding, is also required for the normal longevity responses to both cold and warm temperatures, indicating a potential role of proteostasis in these responses (*14*). Several factors involved in lipid metabolism pathways, including MDT-15/Mediator 15 (*15*), PAQR-2 (*16*), and prostaglandin (*17*), also modulate longevity in response to low temperatures. Measuring the lifespans of a diverse array of short- or long-lived mutant worms at different temperatures revealed that temperature-induced lifespan changes are genetically controlled by temperature-specific gene regulation (*18*). These studies suggest that the longevity response to temperature is a complex and highly regulated process, and that integrative research is needed to define the relationships between temperature, aging mechanisms, and longevity.

Here, using the *C. elegans* model, we demonstrate that functional loss of NPR-8, a neuronal G protein-coupled receptor (GPCR) related to mammalian neuropeptide Y receptors, extends worm lifespan at 25°C but not at 20°C or 15°C, and that the extension in lifespan at 25°C is regulated by the NPR-8-expressing amphid chemosensory neurons AWB and AWC as well as AFD thermosensory neurons, indicating that the nervous system governs longevity in response to higher temperature. Integrative RNA sequencing analysis to examine the relationships between temperature, aging, and NPR-8 revealed that both warm temperature and old age profoundly alter gene expression. Genetic manipulation and functional assays further uncovered that NPR-8-regulated collagen expression contributes to the extended lifespan of *npr-8* mutant animals at 25°C and is also required for maintaining the lifespan of wild-type animals. Since elevated collagen expression is a common feature of many lifespan-extending interventions and enhanced stress resistance to various environmental stimuli (*19, 20*), collagen expression could promote healthy aging and may hold the key to our quest for long and healthy life.

## Results

### Functional loss of NPR-8 extends *C. elegans* lifespan at warm temperature

Three temperatures (15°C, 20°C, and 25°C) are routinely used for *C. elegans* propagation in the laboratory, and their inverse effects on lifespan have been well documented, (i.e., an approximately 75% increase in lifespan occurs with a 5°C drop in temperature) (*4*). During our work on neural regulation of *C. elegans* defense against pathogen infection (*20*), we unexpectedly observed that under noninfectious conditions, *C. elegans* lacking functional NPR-8 (*npr-8(ok1439)* animals) lived longer than wild-type *N2* animals at 25°C, whereas the mutant and wild-type animals had the same lifespan at 20°C or 15°C (Fig. 1A, 1B and 1C). To determine whether the observed longevity phenotype at the warm temperature of 25°C was due to the deletion mutation in *npr-8*, we performed lifespan assays at 25°C using the transgenic strain JRS17 in which *npr-8* is re-expressed under its native promoter in *npr-8(ok1439)* animals (*20*). Re-expression of NPR-8 significantly decreased the extended lifespan of the *npr-8* deletion mutants (Fig. 1D), confirming that the mutation in *npr-8* is indeed responsible for the extended longevity phenotype at warm temperature. To investigate whether this phenotype is allele-specific, we performed lifespan assays using another *npr-8* null strain (*npr-8(ok1446)* animals) that contains a deletion mutation in the *npr-8* gene at a different location than the one in *npr-8(ok1439)* animals (*21*). Our results showed that, like *npr-8(ok1439)* animals, *npr-8(ok1446)* animals also lived longer than wild-type animals at 25°C but had wild-type lifespan at 20°C and 15°C (Fig. S1), indicating that the extended lifespan phenotype is not allele-specific but is caused by the general lack of NPR-8 function. These results suggest that NPR-8 suppresses lifespan at warm temperature, contributing to the inhibitory effect of warm temperature on longevity.

**Fig. 1.**
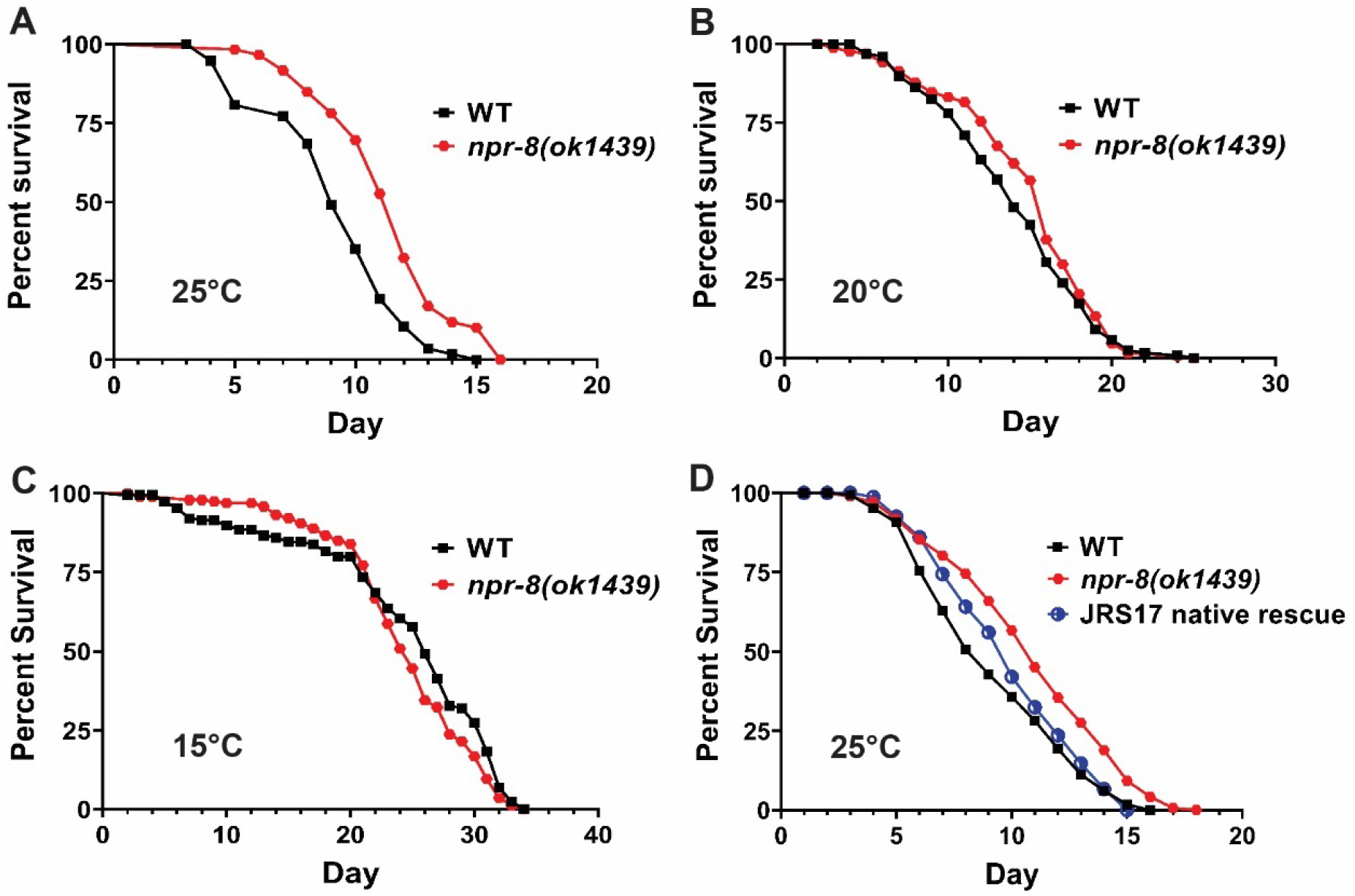
Functional loss of NPR-8 extends lifespan at 25°C but not at 20°C or 15°C. WT and *npr-8(ok1439)* animals were grown on *E. coli* strain OP50 at 25°C **(A)**, 20°C **(B)**, or 15°C **(C)**, and scored for survival over time. The graphs are the combined results of three independent experiments. Each experiment included *n* = 60 adult animals per strain. *p*-values represent the significance level of the mutants relative to the WT, *p* <0.0001 in (A), *p* = 0.1640 in (B), and *p* = 0.1126 in (C). **(D)** WT, *npr-8(ok1439)*, JRS17 (*npr-8* expression restored in *npr-8(ok1439)* under its native promoter) animals were grown on *E. coli* strain OP50 at 25°C and scored for survival over time. The graphs are the combined results of three independent experiments. Each experiment included *n* = 60 adult animals per strain. *p*-values are relative to *npr-8(ok1439)*: WT, *p* < 0.0001; JRS17, *p* < 0.0001.

### The NPR-8-dependent longevity response to warm temperature is neurally regulated

NPR-8 is a neuronal GPCR expressed in the amphid sensory neurons AWB, ASJ and AWC (*20*). Our previous study demonstrated that NPR-8 functions in these neurons to regulate *C. elegans* defense against pathogen infection (*20*). To determine whether the NPR-8-dependent longevity response to warm temperature is also neurally controlled, we took advantage of our established rescue strains that specifically express NPR-8 in the individual neurons under cell-specific promoters in *npr-8(ok1439)* animals (*20*). Lifespan assays at 25°C with these animals showed that re-expressing NPR-8 in AWB or AWC neurons rescued the extended lifespan phenotype of the mutants, while re-expression of NPR-8 in ASJ neurons had no effect on the mutants’ lifespan (Fig. 2A). Like *npr-8(ok1439)* animals, all rescue animals exhibited lifespan similar to that of wild-type animals at 20°C (except the ASJ rescue animals, which displayed slightly longer lifespan than wild type) (Fig. S2). These results indicate that the lack of functional NPR-8 in AWB and AWC neurons is responsible for the mutants’ extended lifespan at 25°C. In other words, NPR-8 functions in AWB and AWC neurons to regulate the longevity response to warm temperature.

**Fig. 2.**
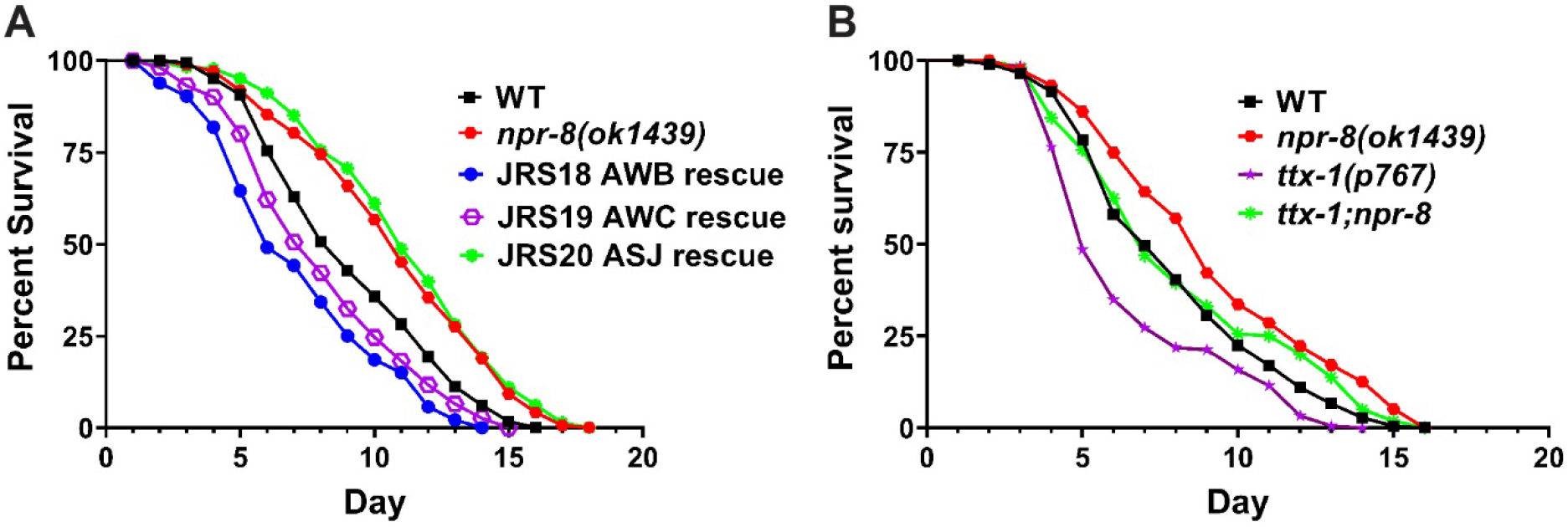
The NPR-8-dependent longevity response to warm temperature is neurally regulated. **(A)** WT, *npr-8 (ok1439)*, and rescue animals were grown at 25°C and scored for survival over time. JRS18, *npr-8* expression rescued in AWB neurons; JRS19, *npr-8* expression rescued in AWC neurons; and JRS20, *npr-8* expression rescued in ASJ neurons. The graphs are the combined results of three independent experiments. Each experiment included *n* = 60 adult animals per strain. *p*-values are relative to *npr-8 (ok1439)*: WT, *p* <0.0001; JRS18, *p* <0.0001; JRS19, *p* <0.0001; JRS20, *p* = 0.3476. **(B)** WT, *npr-8(ok1439), ttx-1(p767), ttx-1;npr-8* animals were grown at 25°C and scored for survival over time. The graphs are the combined results of three independent experiments. Each experiment included *n* = 60 adult animals per strain. *p*-values relative to *npr-8(ok1439)*: WT, *p* = 0.0001; *ttx-1(p767), p* < 0.0001; *ttx-1;npr-8, p* = 0.0101. *p*-values relative to WT: *ttx-1(p767), p* < 0.0001; *ttx-1;npr-8, p* = 0.1872.

In *C. elegans*, AFD thermosensory neurons are required for maintaining lifespan at warm temperature (25°C), and ablation or inactivation of these neurons shortens the worm’s lifespan (*12, 13*). To test whether AFD neurons also function in the NPR-8-dependent longevity response to warm temperature, we examined how a loss-of-function mutation in *ttx-1*, a gene critical for AFD differentiation and function (*22*), affected the lifespan of *npr-8(ok1439)* animals at warm temperature. To this end, we crossed *npr-8(ok1439)* and *ttx-1(p767)* animals to generate *ttx-1;npr-8* double mutant animals, then subjected the single and double mutant animals, along with wild-type controls, to lifespan assays at 25°C. Consistent with previous reports (*12, 13*), the lives of thermosensory-defective *ttx-1(p767)* animals were shorter than those of wild-type animals (Fig. 2B). Interestingly, *ttx-1;npr-8* double mutant animals displayed longer lifespan than *ttx-1(p767)* animals but shorter lifespan than *npr-8(ok1439)* animals (Fig. 2B), indicating that the *ttx-1* and *npr-8* mutations reduce each other’s effect on longevity at warm temperature. These results suggest that the warm temperature-induced longevity response is regulated by a circuit of thermosensory and chemosensory neurons working in an antagonistic manner.

### Both warm temperature and old age broadly alter gene expression in *C. elegans*

To gain molecular insights into the NPR-8-dependent longevity response to temperature, we employed RNA-seq to compare gene expression in young and old wild-type and *npr-8(ok1439)* animals propagated at different temperatures. To this end, we collected five replicates of eight groups of RNA samples (young (0-day-old) and 9-day-old adult wild-type and *npr-8(ok1439)* animals propagated at 20°C and 25°C) (Table 1). These samples were then submitted to the Washington State University Genomics Core for RNA-seq analysis. The resulting sequence data (FASTQ files) were deposited into the National Center for Biotechnology Information (NCBI) Sequence Read Archive (SRA) database through the Gene Expression Omnibus (GEO). The processed gene quantification files and differential expression files were deposited in the GEO. All of these data can be accessed through the GEO with the accession number GSE186202. A summary of our RNA-seq analysis is listed in Table 2.

**Table 1.** Grouping of RNA-seq samples.

| Sample Group | Strain | Propagation Temperature | Adult Age |
| --- | --- | --- | --- |
| 1 | *N2* | 20<sup>0</sup>C | young adult (0-day) |
| 2 | *N2* | 20<sup>0</sup>C | 9-day |
| 3 | *N2* | 25<sup>0</sup>C | 0-day |
| 4 | *N2* | 25<sup>0</sup>C | 9-day |
| 5 | *npr-8(ok1439)* | 20<sup>0</sup>C | 0-day |
| 6 | *npr-8(ok1439)* | 20<sup>0</sup>C | 9-day |
| 7 | *npr-8(ok1439)* | 25<sup>0</sup>C | 0-day |
| 8 | *npr-8(ok1439)* | 25<sup>0</sup>C | 9-day |

**Table 2.** Summary of RNA-seq analysis.

| Influencing factor | Comparison | Sample group | Number of regulated genes | Gene ontology (GO) analysis |  | Data location |
| --- | --- | --- | --- | --- | --- | --- |
| Effects of temperature on gene expression | 0-day-old wild-type animals at 25°C relative to 20°C | Group 3 vs Group 1 | 2,265 genes upregulated | 340 enriched biological processes | cell cycle, reproduction, metabolic/biosynthetic processes, organelle organization, development, DNA replication/repair/transcription/recombination, and other processes | Table S1 |
|  |  |  | 5,069 genes downregulated | 68 attenuated biological processes | signaling, phosphorylation/dephosphorylation, immune response, ion/transmembrane transport, and other processes | Table S2<br>Table S3 |
| Effects of aging on gene expression | wild-type animals at 20°C day 9 relative to day 0 | Group 2 vs Group 1 | 3,792 genes upregulated | 428 enriched biological processes | cell cycle, reproduction/development, metabolism, organelle organization, DNA replication/repair/transcription, and other processes | Table S4 |
|  |  |  | 4,937 genes downregulated | 56 attenuated biological processes | ion/transmembrane transport, phosphorylation/dephosphorylation, signaling, immune response, and other processes | Table S5 |
|  |  |  |  | 65 attenuated molecular functions | "structural constituent of cuticle" is the most significantly reduced | Table S6 |
| Effects of *npr-8* mutation on gene expression | 0-day-old *npr-8(ok1439)* animals at 20°C relative to wild-type controls | Group 5 vs Group 1 | 643 genes upregulated | 7 enriched molecular functions | "structural constituent of cuticle" is the most significantly enriched | Table S7 |
|  |  |  | 272 genes downregulated | No attenuated GO terms |  |  |
|  | 9-day-old *npr-8(ok1439)* animals at 20°C relative to wild-type controls | Group 6 vs Group 2 | 1,487 genes upregulated | 33 enriched biological processes | phosphorylation/dephosphorylation, metabolic process | Table S8 |
|  |  |  | 1,305 genes downregulated | 13 attenuated molecular functions | "structural constituent of cuticle" is the most significantly reduced | Table 3 |
|  | 0-day-old *npr-8(ok1439)* animals at 25°C relative to wild-type controls | Group 7 vs Group 3 | 1,482 genes upregulated | 17 enriched molecular functions | "structural constituent of cuticle" is the most significantly enriched | Table S9 |
|  |  |  | 419 genes downregulated | No attenuated GO terms |  |  |
|  | 9-day-old *npr-8(ok1439)* animals at 25°C relative to wild-type controls | Group 8 vs Group 4 | 1,307 genes upregulated | 29 enriched biological processes | phosphorylation/dephosphorylation, metabolic process | Table S10 |
|  |  |  | 631 genes downregulated | No attenuated GO terms |  |  |

Using the RNA-seq data, we first assessed how propagation temperature affects gene expression by comparing the expression profiles of wild-type young-adult (0-day-old) animals grown at 25°C with those from animals grown at 20°C (Group 3 vs. Group 1 in Table 1). In total, 16,135 genes were identified and quantified with a false discovery rate (FDR) of 5%. Among these genes, 2,265 were upregulated and 5,069 were downregulated at least two-fold in animals grown at 25°C relative to those grown at 20°C. The changes in expression of almost half of the total identified genes (45%) indicate that growth temperature profoundly affects gene expression in *C. elegans*. This might explain, at the molecular level, why temperature is so crucial for life in that even modest variations in temperature can alter many aspects of behavior and physiology including aging (*1*). Gene ontology (GO) analysis of the 2,265 upregulated genes identified 340 significantly enriched biological processes, which can be loosely divided into seven categories: cell cycle, reproduction, metabolic/biosynthetic processes, organelle organization, development, DNA replication/repair/transcription/recombination, and other processes (Table S1). GO analysis of the 5,069 downregulated genes identified 68 significantly attenuated biological processes, which can be loosely divided into five categories: signaling, phosphorylation/dephosphorylation, immune response, ion/transmembrane transport, and other processes (Table S2). It is not clear which regulated biological processes are responsible for the shortened lifespan of *C. elegans* at warm temperature. The enrichment of genes involved in metabolic/biosynthetic processes may reflect an increased rate of metabolism at 25°C relative to 20°C (Table S1), which could provide a molecular basis for the rate of living theory. It is worth noting that among all of the downregulated biological processes (Table S2), signaling is the most significantly reduced and includes reduction in GPCR signaling pathway (GO term GO:0007186), neuropeptide signaling pathway (GO:0007218), synaptic signaling (GO:0099536), nervous system process (GO:0050877), and other signaling processes. A total of 523 genes encoding signaling molecules such as GPCRs and neuropeptides as well as enzymes involved in the metabolism of neurotransmitters or neurohormones contribute to the downregulation of signaling processes (Table S3). These data demonstrate that signaling processes, especially those controlled by the nervous system, are reduced in animals propagated at 25°C compared to those propagated at 20°C, indicating a potential connection between neural signaling and longevity in *C. elegans*. Indeed, signaling from AFD thermosensory neurons and their downstream AIY interneurons are required for maintaining lifespan at 25°C (*12, 13*). However, detailed information about neural circuits that control the longevity response to temperature is lacking.

We next examined how aging affects gene expression by comparing the expression profiles of 9-day-old wild-type adult animals with those of 0-day-old adult animals grown at 20°C (Group 2 vs. Group 1 in Table 1). We found that 3,792 genes were upregulated, and 4,937 genes were downregulated at least 2-fold in old animals relative to young adults. The changes in expression of more than half of the total identified genes indicate that, like temperature, age also has profound effects on *C. elegans* behavior and physiology. GO analysis of the 3,792 upregulated genes identified 428 significantly enriched biological processes, which can be loosely divided into six categories: cell cycle, reproduction/development, metabolism, organelle organization, DNA replication/repair/transcription, and other processes (Table S4). GO analysis of the 4,937 downregulated genes identified 56 significantly attenuated biological processes, which can be loosely divided into five categories: ion/transmembrane transport, phosphorylation/dephosphorylation, signaling, immune response, and other processes (Table S5). These data show that similar biological processes are regulated by old age and warm temperature, suggesting that both conditions could be stressful to *C. elegans*. Interestingly and importantly, GO analysis of the downregulated genes also revealed 65 attenuated molecular functions, with “structural constituent of cuticle” being the most significantly reduced (Table S6A). One hundred and seven downregulated genes contribute to the reduction in cuticle structure activity, among which 88 encode cuticular collagens (Table S6B), indicating that cuticular collagens could be essential for aging and longevity. Indeed, previous studies have shown that the expression levels of collagen genes decline with age and that increased collagen expression is a common feature of multiple conserved longevity pathways and essentially every longevity intervention (*19, 23*). Cuticular collagens have also been implicated in lengthening *C. elegans* lifespan under stress such as pathogen infection and oxidative stress (*19, 20*).

### NPR-8 differentially regulates collagen genes in an age- and temperature-dependent manner

To assess the role of NPR-8 in the temperature-induced longevity response, we compared the expression profiles of young and old *npr-8(ok1439)* animals with those of wild-type animals maintained at different propagation temperatures. For 0-day-old young adults grown at 20°C, 643 genes were upregulated and 272 genes were downregulated at least 2-fold in *npr-8(ok1439)* animals relative to wild-type controls. While GO analysis of the 272 downregulated genes did not yield significantly attenuated biological processes or molecular functions, a similar analysis of the 643 upregulated genes identified seven significantly enriched molecular functions, with the GO term “structural constituent of cuticle” being the most significantly enriched (Table S7A). Sixteen upregulated genes were related to cuticle structure activity, nine of which encoded cuticular collagens (Table S7B). These data indicate that at the normal growth temperature of 20°C, NPR-8 suppresses the expression of a subset of cuticular collagen genes in young-adult wild-type animals. This result is consistent with our previous study showing that NPR-8 suppresses the basal expression of collagen genes (*20*). For 9-day-old adult animals grown at 20°C, 1,487 genes were upregulated and 1,305 genes were downregulated at least 2-fold in *npr-8(ok1439)* animals relative to wild-type controls. The total number of regulated genes tripled at old age compared to young age, indicating that the effects of NPR-8 on *C. elegans* physiology are amplified by aging. GO analysis of the 1,487 upregulated genes identified 33 significantly enriched biological processes, which can be divided into three categories: phosphorylation/dephosphorylation, metabolic process, and other processes (Table S8). GO analysis of the 1,305 downregulated genes revealed 13 attenuated molecular functions, with the GO term “structural constituent of cuticle” being the most significantly reduced (Table 3A). Seventy-three downregulated genes contributed to the decrease in cuticle structure activity, 60 of which encoded cuticular collagens (Table 3B). These data indicate that at old age, NPR-8 upregulates the expression of cuticular collagens in wild-type animals, which is surprising and is in direct contrast with NPR-8’s function at young age when it suppresses cuticular collagens (Table S7). More surprisingly, eight of the nine collagen genes (except *col-98*) that changed expression at young age were also regulated at old age (compare Table 3B to Table S7B). These results are unexpected and raised a perplexing question, that is, how does NPR-8, a single type of GPCR, exerts opposite effects on collagen expression at different stages of the *C. elegans* life cycle? Does *npr-8* influence both fitness and aging similar to genes described in George Williams’ antagonistic pleiotropic theory that have beneficial effects on fitness in early life but are detrimental in later life (*24*)?

**Table 3.**
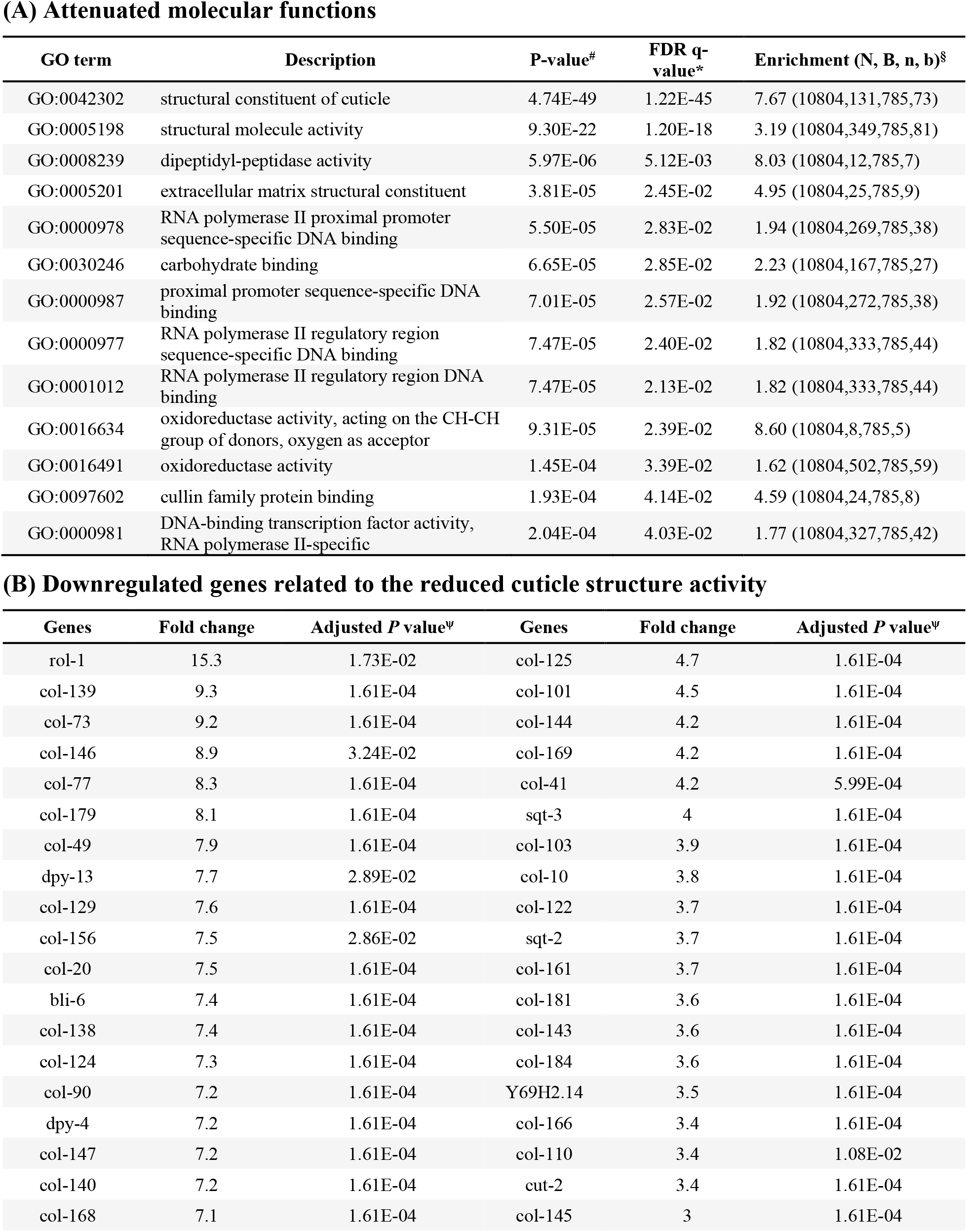
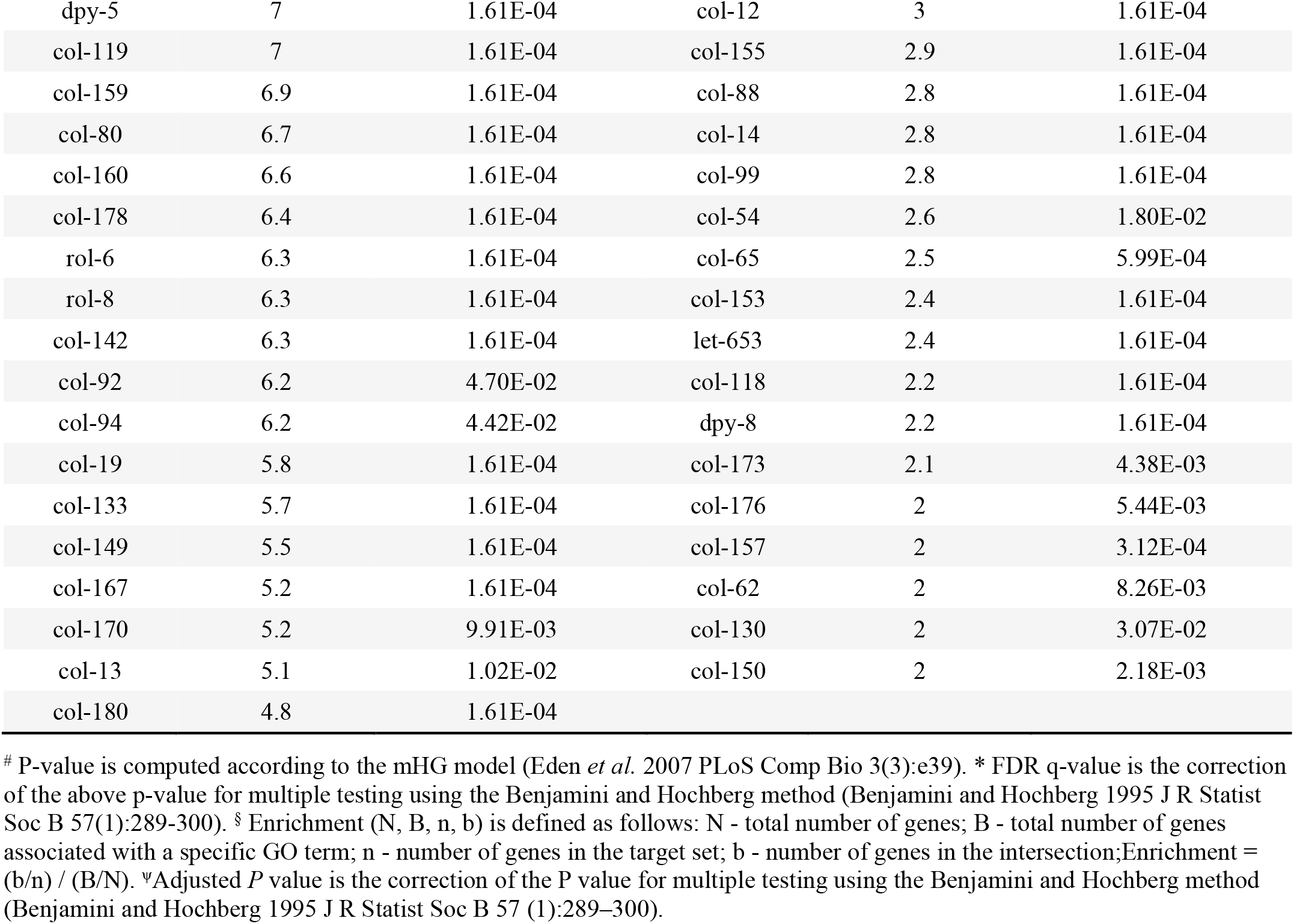
GO analysis of downregulated genes in 9-day-old adult *npr-8(ok1439)* animals grown at 20°C relative to wild-type control animals.

**(A) Attenuated molecular functions**
| GO term | Description | P-value<sup>#</sup> | FDR q-value<sup>*</sup> | Enrichment (N, B, n, b)<sup>§</sup> |
| --- | --- | --- | --- | --- |
| GO:0042302 | structural constituent of cuticle | 4.74E-49 | 1.22E-45 | 7.67 (10804,131,785,73) |
| GO:0005198 | structural molecule activity | 9.30E-22 | 1.20E-18 | 3.19 (10804,349,785,81) |
| GO:0008239 | dipeptidyl-peptidase activity | 5.97E-06 | 5.12E-03 | 8.03 (10804,12,785,7) |
| GO:0005201 | extracellular matrix structural constituent | 3.81E-05 | 2.45E-02 | 4.95 (10804,25,785,9) |
| GO:0000978 | RNA polymerase II proximal promoter sequence-specific DNA binding | 5.50E-05 | 2.83E-02 | 1.94 (10804,269,785,38) |
| GO:0030246 | carbohydrate binding | 6.65E-05 | 2.85E-02 | 2.23 (10804,167,785,27) |
| GO:0000987 | proximal promoter sequence-specific DNA binding | 7.01E-05 | 2.57E-02 | 1.92 (10804,272,785,38) |
| GO:0000977 | RNA polymerase II regulatory region sequence-specific DNA binding | 7.47E-05 | 2.40E-02 | 1.82 (10804,333,785,44) |
| GO:0001012 | RNA polymerase II regulatory region DNA binding | 7.47E-05 | 2.13E-02 | 1.82 (10804,333,785,44) |
| GO:0016634 | oxidoreductase activity, acting on the CH-CH group of donors, oxygen as acceptor | 9.31E-05 | 2.39E-02 | 8.60 (10804,8,785,5) |
| GO:0016491 | oxidoreductase activity | 1.45E-04 | 3.39E-02 | 1.62 (10804,502,785,59) |
| GO:0097602 | cullin family protein binding | 1.93E-04 | 4.14E-02 | 4.59 (10804,24,785,8) |
| GO:0000981 | DNA-binding transcription factor activity, RNA polymerase II-specific | 2.04E-04 | 4.03E-02 | 1.77 (10804,327,785,42) |

**(B) Downregulated genes related to the reduced cuticle structure activity**
| Genes | Fold change | Adjusted *P* value<sup>ψ</sup> | Genes | Fold change | Adjusted *P* value<sup>ψ</sup> |
| --- | --- | --- | --- | --- | --- |
| rol-1 | 15.3 | 1.73E-02 | col-125 | 4.7 | 1.61E-04 |
| col-139 | 9.3 | 1.61E-04 | col-101 | 4.5 | 1.61E-04 |
| col-73 | 9.2 | 1.61E-04 | col-144 | 4.2 | 1.61E-04 |
| col-146 | 8.9 | 3.24E-02 | col-169 | 4.2 | 1.61E-04 |
| col-77 | 8.3 | 1.61E-04 | col-41 | 4.2 | 5.99E-04 |
| col-179 | 8.1 | 1.61E-04 | sqt-3 | 4 | 1.61E-04 |
| col-49 | 7.9 | 1.61E-04 | col-103 | 3.9 | 1.61E-04 |
| dpy-13 | 7.7 | 2.89E-02 | col-10 | 3.8 | 1.61E-04 |
| col-129 | 7.6 | 1.61E-04 | col-122 | 3.7 | 1.61E-04 |
| col-156 | 7.5 | 2.86E-02 | sqt-2 | 3.7 | 1.61E-04 |
| col-20 | 7.5 | 1.61E-04 | col-161 | 3.7 | 1.61E-04 |
| bli-6 | 7.4 | 1.61E-04 | col-181 | 3.6 | 1.61E-04 |
| col-138 | 7.4 | 1.61E-04 | col-143 | 3.6 | 1.61E-04 |
| col-124 | 7.3 | 1.61E-04 | col-184 | 3.6 | 1.61E-04 |
| col-90 | 7.2 | 1.61E-04 | Y69H2.14 | 3.5 | 1.61E-04 |
| dpy-4 | 7.2 | 1.61E-04 | col-166 | 3.4 | 1.61E-04 |
| col-147 | 7.2 | 1.61E-04 | col-110 | 3.4 | 1.08E-02 |
| col-140 | 7.2 | 1.61E-04 | cut-2 | 3.4 | 1.61E-04 |
| col-168 | 7.1 | 1.61E-04 | col-145 | 3 | 1.61E-04 |
| dpy-5 | 7 | 1.61E-04 | col-12 | 3 | 1.61E-04 |
| col-119 | 7 | 1.61E-04 | col-155 | 2.9 | 1.61E-04 |
| col-159 | 6.9 | 1.61E-04 | col-88 | 2.8 | 1.61E-04 |
| col-80 | 6.7 | 1.61E-04 | col-14 | 2.8 | 1.61E-04 |
| col-160 | 6.6 | 1.61E-04 | col-99 | 2.8 | 1.61E-04 |
| col-178 | 6.4 | 1.61E-04 | col-54 | 2.6 | 1.80E-02 |
| rol-6 | 6.3 | 1.61E-04 | col-65 | 2.5 | 5.99E-04 |
| rol-8 | 6.3 | 1.61E-04 | col-153 | 2.4 | 1.61E-04 |
| col-142 | 6.3 | 1.61E-04 | let-653 | 2.4 | 1.61E-04 |
| col-92 | 6.2 | 4.70E-02 | col-118 | 2.2 | 1.61E-04 |
| col-94 | 6.2 | 4.42E-02 | dpy-8 | 2.2 | 1.61E-04 |
| col-19 | 5.8 | 1.61E-04 | col-173 | 2.1 | 4.38E-03 |
| col-133 | 5.7 | 1.61E-04 | col-176 | 2 | 5.44E-03 |
| col-149 | 5.5 | 1.61E-04 | col-157 | 2 | 3.12E-04 |
| col-167 | 5.2 | 1.61E-04 | col-62 | 2 | 8.26E-03 |
| col-170 | 5.2 | 9.91E-03 | col-130 | 2 | 3.07E-02 |
| col-13 | 5.1 | 1.02E-02 | col-150 | 2 | 2.18E-03 |
| col-180 | 4.8 | 1.61E-04 |  |  |  |

We next examined the gene expression profiles of young and old *npr-8(ok1439)* and wild-type animals grown at 25°C. For 0-day-old young adults, 1,482 genes were upregulated and 419 genes were downregulated at least 2-fold in *npr-8(ok1439)* animals relative to wild-type animals. GO analysis of the 1,482 upregulated genes identified 17 enriched molecular functions, with the GO term “structural constituent of cuticle” being the most significantly enriched (Table S9A). Seventy upregulated genes were related to cuticle structure activity, 56 of which encoded cuticular collagens (Table S9B). These data indicate that, similar to 0-day-old young adults grown at 20°C (Table S7), NPR-8 also suppresses the expression of cuticular collagen genes in young adults grown at 25°C. However, there are many more collagen genes suppressed at 25°C than at 20°C (56 genes vs. 9 genes, compare Table S9B with Table S7B), indicating that warm temperature broadens NPR-8’s inhibitory effect on collagen gene expression in young animals. For 9-day-old adults, 1,307 genes were upregulated and 631 genes were downregulated at least 2-fold in *npr-8(ok1439)* animals relative to wild-type animals. GO analysis of the 1,307 upregulated genes revealed 29 significantly enriched biological processes, which can be divided into three categories: phosphorylation/dephosphorylation, metabolic process, and other processes (Table S10). These data demonstrate that in old animals, the biological processes suppressed by NPR-8 at 25°C are similar to those suppressed by NPR-8 at 20°C (compare Table S10 with Table S8), indicating that these specific NPR-8 effects are age-related, not temperature-dependent. GO analysis of the 631 downregulated genes did not yield any significantly enriched biological processes or molecular functions. This is in striking contrast with old worms grown at 20°C in which a large number of cuticular collagen genes (60 genes) were downregulated in *npr-8(ok1439)* animals relative to wild-type animals (Table 3B). Because collagen expression promotes longevity (*19, 23*), this lack of downregulation of collagen expression at 25°C may explain the extended lifespan of *npr-8(ok1439)* animals at warm temperature.

### NPR-8-regulated collagen genes contribute to the extended lifespan of *npr-8(ok1439)* animals at 25°C

To determine whether collagen expression plays a role in *npr-8(ok1439)* animals’ extended lifespan at warm temperature, we inactivated NPR-8-regulated collagen genes using RNA interference (RNAi) or deletion mutation in *npr-8(ok1439)* and wild-type animals and then performed lifespan assays at 25°C on these animals. For the five most-regulated collagen genes tested (downregulated most in *npr-8(ok1439)* animals at 20°C but lacking such downregulation at 25°C, Table 3B), inactivation of each of them (*rol-1, col-49, co-77, col-139*, or *col-179*) significantly suppressed the extended lifespan of *npr-8* mutant animals at 25°C (Fig. 3A and 3C). Inactivation of *col-49* and *col-179* also suppressed the lifespan of wild-type animals (Fig. 3B). These results indicate that the expression of NPR-8-regulated collagen genes contributes to the extended lifespan of *npr-8(ok1439)* animals at 25°C and is also essential for maintaining the lifespan of wild-type animals at warm temperature. This is consistent with the longevity-promoting role of collagens in the aging process (*19, 23*).

**Fig. 3.**
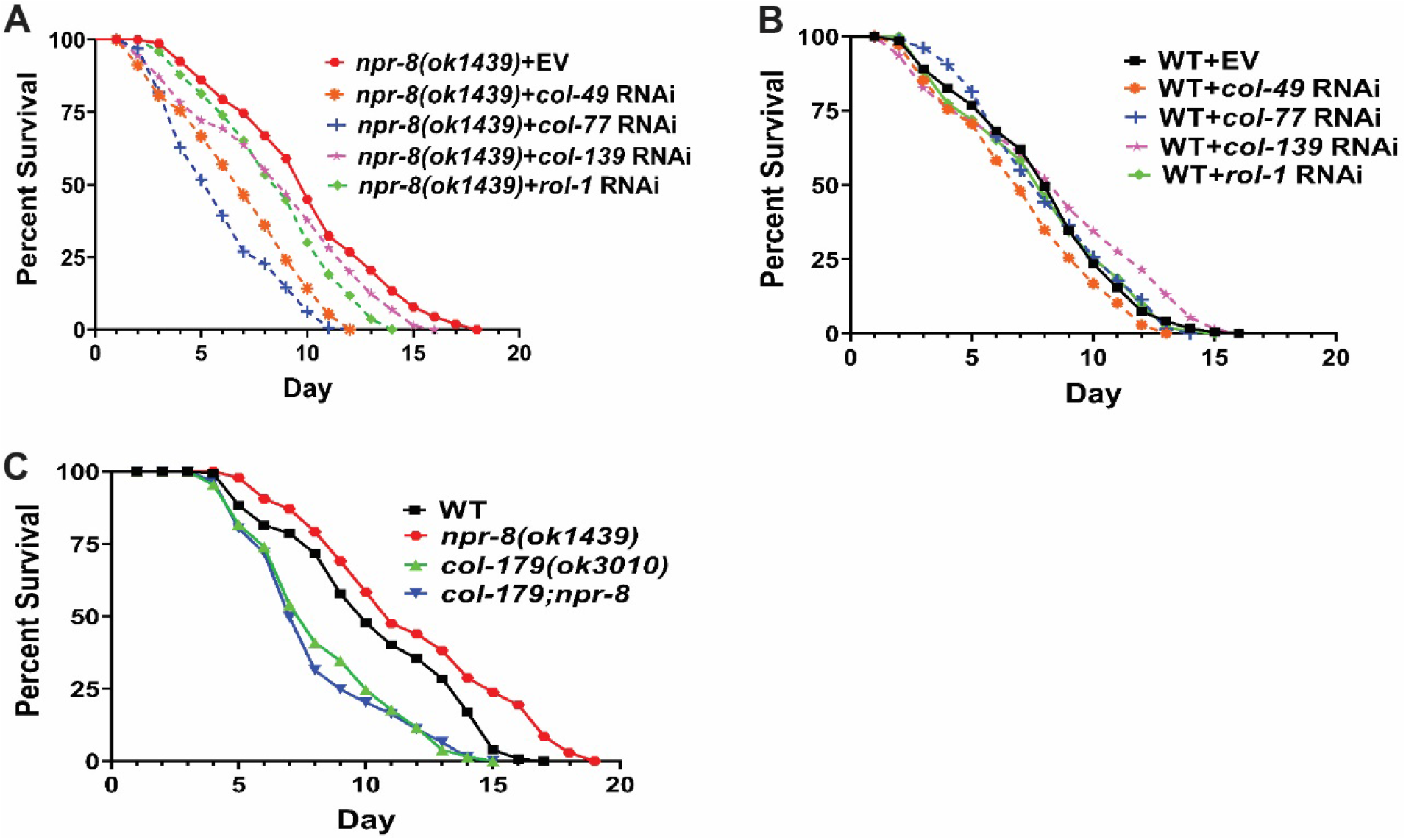
NPR-8-regulated collagen genes are required for warm temperature-induced lifespan extension in *npr-8(ok1439)* animals. *npr-8(ok1439)* animals **(A)** and WT animals **(B)** were grown on dsRNA for empty vector (EV) or for *rol-1, col-49, col-77*, or *col-139* at 25°C and scored for survival over time. The graphs are the combined results of three independent experiments. Each experiment included *n* = 60 adult animals per strain. *p*-values are relative to *npr-8(ok1439)*+EV in (A): *npr-8(ok1439)*+*col-49, p* <0.0001; *npr-8(ok1439)*+*col-77, p* <0.0001; *npr-8(ok1439)*+*col-139, p* = 0.0009; *npr-8 (ok1439)*+*rol-1, p* <0.0001. *p*-values are relative to WT+EV in (B): WT+*col-49, p* = 0.0039; WT+*col-77, p* = 0.9225; WT+*col-139, p* =0.0261; WT+*rol-1, p* = 0.8383. **(C)** WT, *npr-8(ok1439), col-179(ok3010), npr-8;col-179* animals were grown at 25°C and scored for survival over time. The graphs are the combined results of three independent experiments. Each experiment included *n* = 60 adult animals per strain. *p-*values relative to *npr-8(ok1439)*: WT, *p* < 0.0001; *col-179(ok3010), p* < 0.0001; *npr-8;col-179, p* < 0.0001. *p-*values relative to WT: *col-179(ok3010), p* < 0.0001; *npr-8;col-179, p* < 0.0001.

### *npr-8(ok1439)* animals have a less-wrinkled cuticle and appear younger than age-matched wild-type animals

During our study, we observed that *npr-8* mutant animals appeared categorically younger than age-matched wild-type animals by exhibiting fewer wrinkles in the cuticle. To investigate this phenomenon in detail, we performed scanning electron microscopy (SEM) to examine the cuticle of wild-type and *npr-8(ok1439)* animals at various ages (3-, 6-, and 9-day old adults) under the two different propagation temperatures of 20°C and 25°C. Characteristic structural features of the *C. elegans* cuticle include small circumferential furrows separated by broader ridges called annuli and longitudinally oriented ridges called alae (Fig. 4A) (*25*). Distinct alae are considered an indicator of worm adulthood (*26*). Indeed, we observed that 3-day-old and 6-day-old wild-type adult animals propagated at 20°C exhibited smooth cuticle with distinct alae structures (Fig. 4A and 4B). At the same temperature, 9-day-old wild-type adults displayed a wrinkled cuticle with distorted alae (Fig. 4C), whereas 9-day-old *npr-8(ok1439)* animals maintained distinct alae structures on a smooth surface (Fig. 4I). These results demonstrated that there are NPR-8-dependent differences in the cuticles of mutant and wild-type animals. In contrast to what was observed at 20°C, wrinkled cuticles and disintegrated alae were observed among 6-day-old wild-type adults at 25°C (Fig. 4E), and by day 9, alae were disrupted (Fig. 4F), suggesting that warm temperature speeds up aging. Compared to wild-type adults, both 6-day-old and 9-day-old *npr-8(ok1439)* animals displayed a smooth cuticle with distinct structural morphology at 25°C (Fig. 4K and 4L), indicating younger-than-wild-type physiological age and perhaps better overall health. Why would the functional loss of a neuronal GPCR lead to a less-wrinkled cuticle? Since the 1930s, researchers have known that in humans, a finger with nerve damage will not wrinkle under water (*27*). In fact, a medical test, the Wrinkling Test, has been developed to check the wrinkling of patients to assess their possible peripheral nerve damage (*28*). However, the molecular connections between wrinkles and the nerve damage remain unknown. Previously, we have shown that NPR-8-regulated collagen genes are expressed in the cuticle as well as in hypodermal and rectal cells (*20*). The difference in cuticle appearance between *npr-8* mutants and wild-type animals could be due to the differential expression of cuticular collagen genes in these animals, as revealed by the above-described RNA-seq data. Different collagen compositions in the skin controlled by the nerve could also explain the Wrinkling Test.

**Fig. 4.**
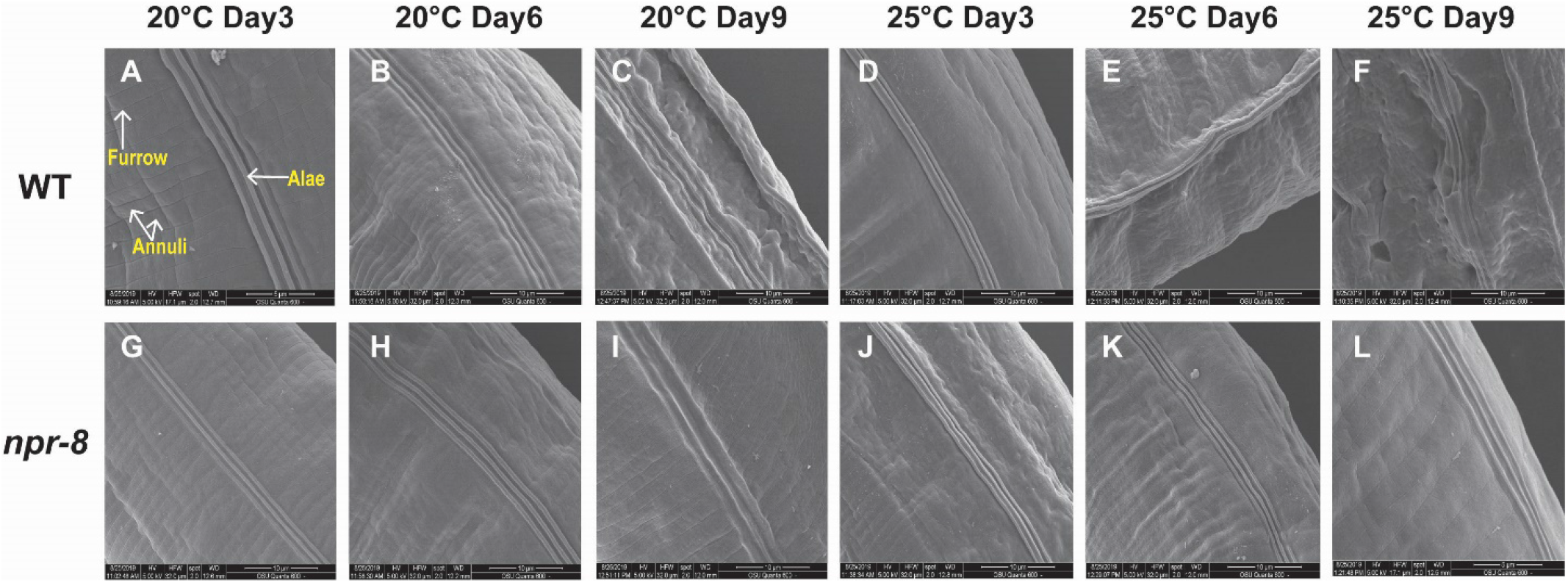
The cuticle of *npr-8(ok1439)* animals was less wrinkled than that of WT animals at old ages at 20°C and 25°C. Adult animals were collected at days 3, 6 and 9 and imaged using FEI Quanta 600 FEG scanning electron microscope.

## Discussion

As in many other invertebrate and vertebrate animals, temperature has inverse effects on longevity in *C. elegans*. Mechanistically, this inverse relationship is not well understood. In the current study, we have demonstrated that functional loss of NPR-8 increases *C. elegans* lifespan at 25°C but not at 20°C and 15°C, indicating that NPR-8 is involved in the longevity response to warm temperature. Further investigation revealed that such regulation is achieved by altering the expression of a subset of collagen genes, and that this regulation is controlled by the NPR-8-expressing amphid chemosensory neurons AWB and AWC as well as AFD thermosensory neurons. These two types of sensory neurons work in an antagonistic manner to maintain longevity at 25°C. Therefore, an NPR-8-dependent neural-longevity regulatory circuit has emerged whereby the nervous system controls the longevity response to warm temperature by regulating collagen expression. However, many details of this regulatory circuit are lacking. For example, which neurons form a network with AWB, AWC and AFD neurons to control longevity at warm temperature? What neuroendocrine signaling pathways are used to relay neural signals to non-neural tissues, since we have previously found that NPR-8-regulated collagens are primarily expressed in the cuticle and hypodermal and rectal cells (*20*)? How does collagen expression influence aging? Addressing these questions will provide important mechanistic insights into the temperature-induced longevity response and define the role of the nervous system in this process.

The inverse relationship between longevity and temperature has long been explained using the rate of living theory, which posits that higher temperatures increase chemical reaction rates, thus speeding up the aging process (*8*). However, recent research has identified specific molecules, cells, and signaling pathways that are involved in the longevity response to temperature, indicating that this response is a regulated process and not simply due to changes in thermodynamics (*1, 9*). The actual mechanisms behind the inverse relationship seem not quite that clear. On the one hand, our study provides further evidence supporting that the temperature-induced longevity response is indeed a regulated process. We have identified the neuronal GPCR NPR-8 and the AWB, AWC and AFD neurons that function in the *C. elegans* longevity response to warm temperature, indicating that this response is controlled by the nervous system. On the other hand, our RNA-seq data revealed the enrichment of many genes involved in metabolic/biosynthetic processes in wild-type animals at 25°C relative to 20°C (Table S1), reflecting increased metabolism at warm temperature. Thus, our data provide a partial molecular basis for the rate of living theory, supporting the idea that there is some scientific truth in this theory. As the rate of living theory and the view of the temperature-induced longevity response being a regulated process are not mutually exclusive, we propose a hybrid hypothesis in which higher temperatures speed up chemical reactions and the aging process in both thermodynamic and neurally regulated manners in living organisms.

An important finding of our current study is that the expression of NPR-8-regulated collagen genes is required for maintaining the lifespan of wild-type animals at 25°C, and that the expression of these genes contributes to the extended lifespan of *npr-8(ok1439)* animals at warm temperature (Fig. 3). Collagen expression has also been implicated in enhanced resistance to pathogen infection (*20*) and oxidative stress (*19*). In fact, elevated collagen expression is a common feature of multiple conserved longevity pathways and essentially every longevity intervention (*19, 29–31*). These findings indicate a general role of collagen in aging. As stress resistance is a critical parameter for assessing healthy lifespan, or healthspan (*32, 33*), collagen expression likely promotes healthy aging. Indeed, we observed that *npr-8* mutant animals with increased collagen expression had a less-wrinkled cuticle and appeared younger than age-matched wild-type animals (Fig. 4). Remarkably, mammalian studies have shown that placing senescent cells or aged stem cells in “young extracellular matrix” (ECM; collagen is the predominant component of ECM) rejuvenates them (*34–36*). How such ECM-cell interactions reverse the cellular aging process remains a mystery. It is reasonable to speculate that the signaling role of collagens, which is less understood than their structural role (*37*), is likely involved in aging. Indeed, collagens have been shown to bind to diverse families of cell-surface receptors, including integrins, receptor tyrosine kinases, and immunoglobulin-like receptors, thus influencing cellular physiology and behavior (*38, 39*). The integrin-signaling complex, acting as a major link between collagen and the cytoskeleton, has been implicated in modulating longevity and thermotolerance (*40*). These studies suggest that research deciphering how collagen expression promotes healthy aging may hold the key to the human request for long and healthy life.

## Materials and Methods

### *C. elegans* strains

*C. elegans* strains were cultured under standard conditions and fed *E. coli* OP50. Wild-type animals were *C. elegans N2* Bristol. *npr-8 (ok1439), npr-8(ok1446), ttx-1 (p767)* (AFD-defective mutant), and *col-179(ok3010)* animals were obtained from the Caenorhabditis Genetics Center (University of Minnesota, Minneapolis, MN) and backcrossed with wild-type N2 for 3 to 8 times. JRS30 [*npr-8(ok1439);col-179(ok3010)*], YYL11 [*ttx-1(p767);npr-8(ok1439)*], JRS17 [*npr-8p(2kb)::npr-8 gDNA::gfp*] (native NPR-8 rescue strain), JRS18 [*str-1p(1*.*3kb)::npr-8 gDNA::gfp*] (NPR-8 rescue in AWB neurons), JRS19 [*str-2p(2kb)::npr-8 gDNA::gfp*] (NPR-8 rescue in AWC neurons), and JRS20 [*trx-1p(1*.*1kb)::npr-8 gDNA::gfp*] (NPR-8 rescue in ASJ neurons) were generated using standard genetic techniques (*20*).

### Lifespan assay

Wild-type and mutant worms were maintained at 20°C and fed *E. coli* OP50 on NGM medium. For temperature-specific assays at 15°C and 25°C, worm strains were maintained for at least two generations at the respective temperature before being used for egg synchronization. For egg synchronization, 3×20 gravid adult worms per strain were used. The time window of egg synchronization was set for 1 hr, during which all animals were maintained at the respective assay temperature (15°C, 20°C, or 25°C). *E. coli OP50* lawn was prepared by spreading a 50-µl drop of an overnight culture of bacteria on the NGM medium (3.5-cm petri plates). Plates were incubated at 37°C for 16 hrs and then cooled down at room temperature for at least 1 hr before seeding with synchronized worms. 3×20 synchronized worms for each group were transferred onto the live *E. coli* OP50 plates. Worms were counted and transferred to fresh, live *E. coli* OP50 culture daily until death. All assays were performed in triplicates for each strain.

### RNA isolation

To gain molecular insights into the NPR-8-dependent longevity response to temperature, we employed RNA-seq to compare gene expression in young and old wild-type and *npr-8(ok1439)* animals propagated at different temperatures. To this end, we collected five replicates of eight groups of RNA samples (young (0-day-old) and 9-day-old adult wild-type and *npr-8(ok1439)* animals propagated at 20°C and 25°C) (Table 1). The temperature-specific worm cultivation and synchronization steps were performed as described in the lifespan assay section. A portion of synchronized adult animals were collected at the age of 0-day old, and the remaining animals were grown till day 9 before collection. These animals were washed every day using M9 buffer and the adult animals were filtered using cell strainer (40-µm nylon filter, Falcon) and transferred to *E. coli* OP50-seeded NGM plates. Worms were washed and collected with M9 buffer then snap-frozen in the TRIzol reagent (Thermo Fisher Scientific) and stored at −80°C until RNA isolation. RNA was extracted using the QIAzol lysis reagent (Qiagen) and purified with the RNeasy Plus Universal Kit (Qiagen).

### RNA sequencing

Total RNA samples were obtained as described above and submitted to the WSU Genomics Core for RNA-seq analysis. RNA-seq and related bioinformatic analyses were performed following our published protocols with modifications (*20*). Specifically, after alignment of FASTQ files to the *C. elegans* reference genome (ce10, UCSC) using HISAT2 (*41*), gene expression quantification and differential expression were analyzed using Cuffquant and Cuffdiff, respectively (*42*). The raw sequence data (FASTQ files) were deposited in the National Center for Biotechnology Information (NCBI) Sequence Read Archive (SRA) database through the Gene Expression Omnibus (GEO). The processed gene quantification files and differential expression files were deposited in the GEO. All of these data can be accessed through the GEO with the accession number GSE186202.

### Scanning electron microscopy (SEM) analysis

To determine age- and temperature-dependent structural changes in the cuticle, we performed SEM to examine wild-type and *npr-8 (ok1439)* animals at different ages (3-, 6-, and 9-day-old adults) propagated at 20°C or 25°C. Temperature-specific worm maintenance and synchronization were the same as described in the lifespan assay. Sample preparation and image acquisition were performed as we previously described (*20*).

### RNA interference

The collagen genes *col-77, col-49, col-139*, and *rol-1* were individually silenced using RNAi in wild-type and *npr-8 (ok1439)* animals at 25°C. The temperature-specific worm cultivation and synchronization steps were performed as described in the above lifespan assay section. The RNAi protocol was performed as we previously described (*20*). The impact of collagen gene silencing on *C. elegans* lifespan was assessed using lifespan assays.

### Statistical analysis

For *C. elegans* lifespan assays, animal survival was plotted as a non-linear regression curve using the PRISM computer program (version 9, GraphPad Software, Inc. La Jolla, CA). Lifespan curves were considered different than the appropriate controls when *p*-values were < 0.05. Prism uses the product limit or Kaplan-Meier method to calculate survival fractions and the logrank test (equivalent to the Mantel-Heanszel test) to compare survival curves. All of the experiments were repeated at least 3 times, unless otherwise indicated.

## Acknowledgements

We thank Dr. Collin Ewald at the Swiss Federal Institute of Technology in Zurich and Dr. Seung-Jae Lee at the Korea Advanced Institute of Science and Technology for critical reading of the manuscript. We thank Dr. Jingru Sun at Washington State University for providing us with experimental reagents, worm strains, and technical support. Some worm strains were provided by the *Caenorhabditis* Genetics Center, which is funded by the NIH Office of Research Infrastructure Programs (P40 OD010440).

## Author contributions

S.N.P., D.S., and Y.L. designed and performed experiments and analyzed data. S.N.P. and Y.L. wrote the paper.

## Data and materials availability

The RNA sequencing data have been deposited in NCBI’s Sequence Read Archive (SRA) database through the Gene Expression Omnibus (GEO). Processed gene quantification files and differential expression files have been deposited in GEO. All of these data can be accessed through the GEO with the accession number GSE186202. All data needed to evaluate the conclusions in the paper are present in the paper and/or the Supplementary Materials. Additional data are available from the authors upon request. The *C. elegans* strains and plasmids constructed by us and the primer sequences used for their construction are available upon request.

## Supplementary Materials

**Table S1. Enriched biological processes in 0-day-old adult wild-type animals at 25°C vs 20°C**

**Table S2. Attenuated biological processes in 0-day-old adult wild-type animals at 25°C vs 20°C**

**Table S3. Downregulated genes contributing to the attenuated signaling processes in 0-day-old adult wild-type animals at 25°C vs 20°C**

**Table S4. Enriched biological processes in 9-day-old wild-type animals at 20°C vs 0-day-old adult animals**

**Table S5. Attenuated biological processes in 9-day-old wild-type animals at 20°C vs 0-day-old adult animals**

**Table S6. Gene ontology (GO) analysis of downregulated genes in 9-day-old wild-type animals grown at 20°C relative to 0-day-old adult animals**

**Table S7. GO analysis of upregulated genes in 0-day-old adult *npr-8(ok1439)* animals grown at 20°C relative to wild-type animals**

**Table S8. Enriched biological processes in 9-day-old *npr-8(ok1439)* animals grown at 20°C vs wild-type animals**

**Table S9. GO analysis of upregulated genes in 0-day-old adult *npr-8(ok1439)* animals grown at 25°C relative to wild-type animals**

**Table S10. Enriched biological processes in 9-day-old *npr-8(ok1439)* animals grown at 25°C vs wild-type animals**

**Fig. S1.**
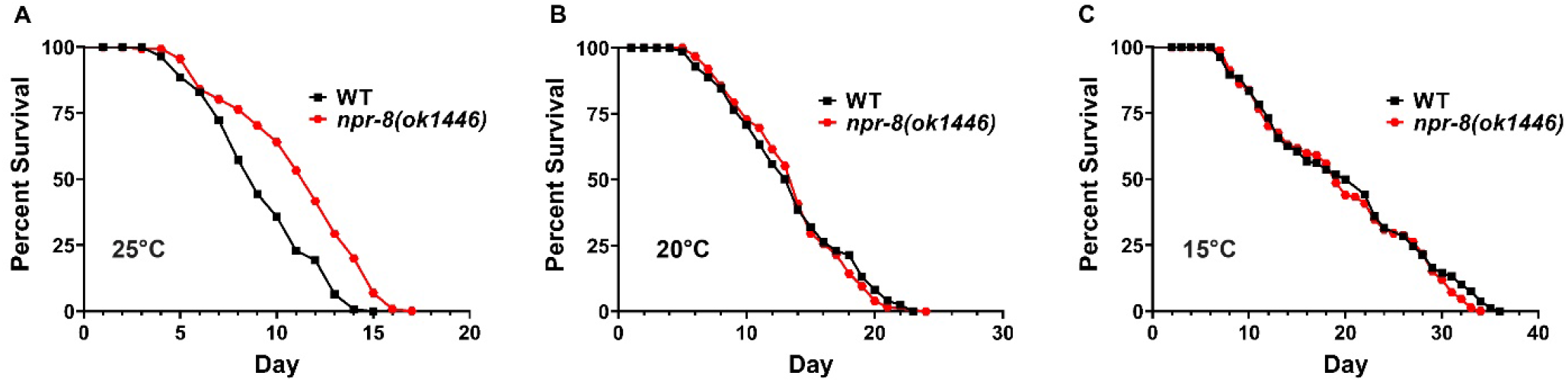
*npr-8(ok1446)* animals live longer than WT animals at 25°C but not at 20°C or 15°C. WT and *npr-8(ok1446)* animals were grown on *E. coli* strain OP50 at 25°C **(A)**, 20°C **(B)**, or 15°C **(C)**, and scored for survival over time. The graphs are the combined results of three independent experiments. Each experiment included *n* = 60 adult animals per strain. *p*-values represent the significance level of the mutants relative to the WT, *p* <0.0001 in (A), *p* = 0.8114 in (B), and *p* = 0.2202 in (C).

**Fig. S2.**
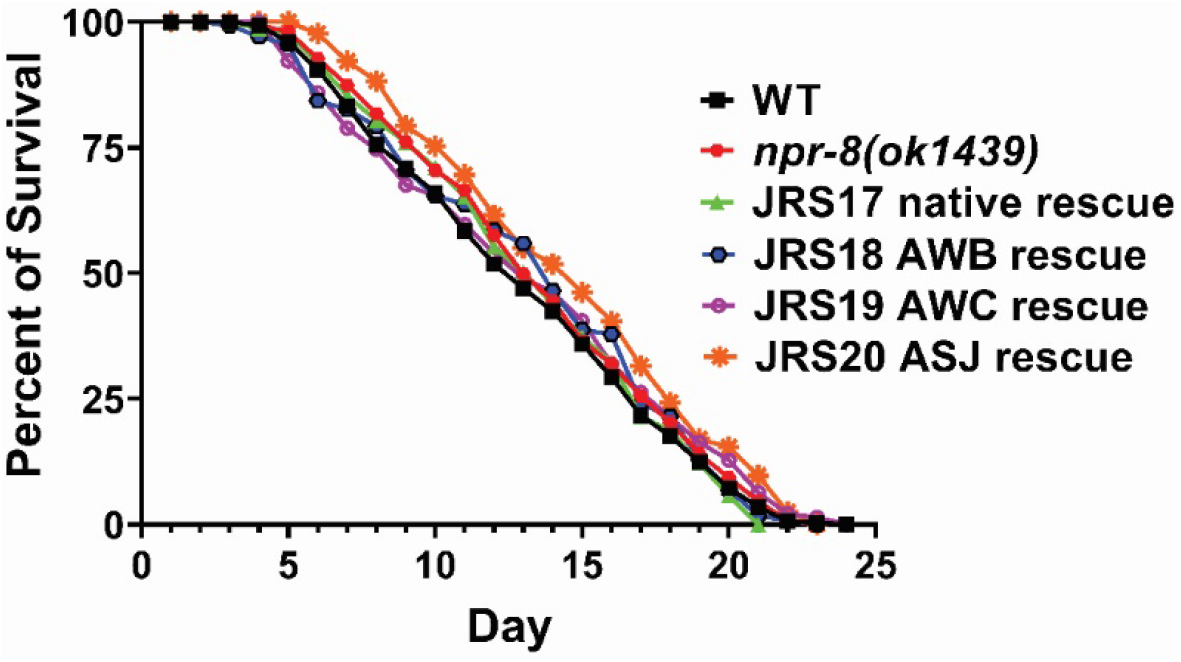
NPR-8 rescue animals exhibited lifespan similar to that of wild-type animals at 20°C. WT, *npr-8 (ok1439)*, and rescue animals were grown at 20°C and scored for survival over time. JRS17, *npr-8* expression restored in *npr-8(ok1439)* under its native promoter; JRS18, *npr-8* expression rescued in AWB neurons; JRS19, *npr-8* expression rescued in AWC neurons; and JRS20, *npr-8* expression rescued in ASJ neurons. The graphs are the combined results of three independent experiments. Each experiment included *n* = 60 adult animals per strain. *p*-values are relative to WT: *npr-8 (ok1439), p* = 0.3424; JRS17, *p* = 0.9600; JRS18, *p* = 0.6348; JRS19, *p* = 0.9119; JRS20, *p* = 0.0066.

**Table S1 – S10**

https://pounce.spokane.wsu.edu/index.php/s/zrDyryXGq32pe85

## References

1. G. Keil, E. Cummings, J. P. de Magalhaes, Being cool: how body temperature influences ageing and longevity. Biogerontology 16, 383–397 (2015).

2. J. Loeb, J. H. Northrop, Is There a Temperature Coefficient for the Duration of Life? Proceedings of the National Academy of Sciences of the United States of America 2, 456–457 (1916).

3. J. W. Macarthur, W. H. Baillie, Metabolic activity and duration of life. I. Influence of temperature on longevity in Daphnia magna. Journal of Experimental Zoology 53, 221–242 (1929).

4. W. A. Van Voorhies, S. Ward, Genetic and environmental conditions that increase longevity in Caenorhabditis elegans decrease metabolic rate. Proceedings of the National Academy of Sciences of the United States of America 96, 11399–11403 (1999).

5. R. L. Walford, R. K. Liu, Husbandry, life span, and growth rate of the annual fish, Cynolebias adloffi E. Ahl. Experimental Gerontology 1, 161–168 (1965).

6. B. Conti, M. Sanchez-Alavez, R. Winsky-Sommerer, M. C. Morale, J. Lucero, S. Brownell, V. Fabre, S. Huitron-Resendiz, S. Henriksen, E. P. Zorrilla, L. de Lecea, T. Bartfai, Transgenic mice with a reduced core body temperature have an increased life span. Science 314, 825–828 (2006).

7. B. A. Rikke, T. E. Johnson, Lower body temperature as a potential mechanism of life extension in homeotherms. Exp Gerontol 39, 927–930 (2004).

8. L. A. Demetrius, Boltzmann, Darwin and Directionality theory. Physics Reports 530, 1–85 (2013).

9. B. Kim, J. Lee, Y. Kim, S. V. Lee, Regulatory systems that mediate the effects of temperature on the lifespan of Caenorhabditis elegans. J Neurogenet 34, 518–526 (2020).

10. R. Xiao, B. Zhang, Y. Dong, J. Gong, T. Xu, J. Liu, X. Z. Xu, A genetic program promotes C. elegans longevity at cold temperatures via a thermosensitive TRP channel. Cell 152, 806–817 (2013).

11. B. Zhang, R. Xiao, E. A. Ronan, Y. He, A. L. Hsu, J. Liu, X. Z. Xu, Environmental Temperature Differentially Modulates C. elegans Longevity through a Thermosensitive TRP Channel. Cell reports 11, 1414–1424 (2015).

12. S. J. Lee, C. Kenyon, Regulation of the longevity response to temperature by thermosensory neurons in Caenorhabditis elegans. Current biology : CB 19, 715–722 (2009).

13. Y. C. Chen, H. J. Chen, W. C. Tseng, J. M. Hsu, T. T. Huang, C. H. Chen, C. L. Pan A C. elegans Thermosensory Circuit Regulates Longevity through crh-1/CREB-Dependent flp-6 Neuropeptide Signaling. Developmental cell 39, 209–223 (2016).

14. M. Horikawa, S. Sural, A. L. Hsu, A. Antebi, Co-chaperone p23 regulates C. elegans Lifespan in Response to Temperature. PLoS genetics 11, e1005023 (2015).

15. D. Lee, S. W. A. An, Y. Jung, Y. Yamaoka, Y. Ryu, G. Y. S. Goh, A. Beigi, J. S. Yang, G. Y. Jung, D. K. Ma, C. M. Ha, S. Taubert, Y. Lee, S. V. Lee, MDT-15/MED15 permits longevity at low temperature via enhancing lipidostasis and proteostasis. PLoS biology 17, e3000415 (2019).

16. Y. L. Chen, J. Tao, P. J. Zhao, W. Tang, J. P. Xu, K. Q. Zhang, C. G. Zou, Adiponectin receptor PAQR-2 signaling senses low temperature to promote C. elegans longevity by regulating autophagy. Nat Commun 10, 2602 (2019).

17. H. J. Lee, A. Noormohammadi, S. Koyuncu, G. Calculli, M. S. Simic, M. Herholz, A. Trifunovic, D. Vilchez, Prostaglandin signals from adult germ stem cells delay somatic aging of Caenorhabditis elegans. Nat Metab 1, 790–810 (2019).

18. H. Miller, M. Fletcher, M. Primitivo, A. Leonard, G. L. Sutphin, N. Rintala, M. Kaeberlein, S. F. Leiser, Genetic interaction with temperature is an important determinant of nematode longevity. Aging Cell 16, 1425–1429 (2017).

19. C. Y. Ewald, J. N. Landis, J. Porter Abate, C. T. Murphy, T. K. Blackwell, Dauer-independent insulin/IGF-1-signalling implicates collagen remodelling in longevity. Nature 519, 97–101 (2015).

20. D. Sellegounder, Y. Liu, P. Wibisono, C. H. Chen, D. Leap, J. Sun, Neuronal GPCR NPR-8 regulates C. elegans defense against pathogen infection. Sci Adv 5, eaaw4717 (2019).

21. C. e. D. M. Consortium, large-scale screening for targeted knockouts in the Caenorhabditis elegans genome. G3 (Bethesda) 2, 1415–1425 (2012).

22. J. S. Satterlee, H. Sasakura, A. Kuhara, M. Berkeley, I. Mori, P. Sengupta, Specification of thermosensory neuron fate in C. elegans requires ttx-1, a homolog of otd/Otx. Neuron 31, 943–956 (2001).

23. Y. V. Budovskaya, K. Wu, L. K. Southworth, M. Jiang, P. Tedesco, T. E. Johnson, S. K. Kim, An elt-3/elt-5/elt-6 GATA transcription circuit guides aging in C. elegans. Cell 134, 291–303 (2008).

24. G. C. Williams, Pleiotropy, natural selection, and the evolution of senescence. Evolution 11, 14 (1957).

25. R. Lints, D. H. Hall, The cuticle. in WormAtlas, (2009).

26. Z. F. Altun, D. H. Hall, Introduction to C. elegans anatomy. In WormAtlas, (2009).

27. S. O’Riain, New and simple test of nerve function in hand. Br Med J 3, 615–616 (1973).

28. T. M. Vasudevan, A. M. van Rij, H. Nukada, P. K. Taylor, Skin wrinkling for the assessment of sympathetic function in the limbs. Aust N Z J Surg 70, 57–59 (2000).

29. J. Myllyharju, K. I. Kivirikko, Collagens, modifying enzymes and their mutations in humans, flies and worms. Trends in genetics : TIG 20, 33–43 (2004).

30. B. H. Toyama, M. W. Hetzer, Protein homeostasis: live long, won’t prosper. Nature reviews. Molecular cell biology 14, 55–61 (2013).

31. J. Varani, M. K. Dame, L. Rittie, S. E. Fligiel, S. Kang, G. J. Fisher, J. J. Voorhees, Decreased collagen production in chronologically aged skin: roles of age-dependent alteration in fibroblast function and defective mechanical stimulation. Am J Pathol 168, 1861–1868 (2006).

32. C. Y. Ewald, J. I. Castillo-Quan, T. K. Blackwell, Untangling Longevity, Dauer, and Healthspan in Caenorhabditis elegans Insulin/IGF-1-Signalling. Gerontology 64, 96–104 (2018).

33. A. Bansal, L. J. Zhu, K. Yen, H. A. Tissenbaum, Uncoupling lifespan and healthspan in Caenorhabditis elegans longevity mutants. Proceedings of the National Academy of Sciences of the United States of America 112, E277–286 (2015).

34. H. R. Choi, K. A. Cho, H. T. Kang, J. B. Lee, M. Kaeberlein, Y. Suh, I. K. Chung, S. C. Park, Restoration of senescent human diploid fibroblasts by modulation of the extracellular matrix. Aging Cell 10, 148–157 (2011).

35. Y. Sun, W. Li, Z. Lu, R. Chen, J. Ling, Q. Ran, R. L. Jilka, X. D. Chen, Rescuing replication and osteogenesis of aged mesenchymal stem cells by exposure to a young extracellular matrix. FASEB J 25, 1474–1485 (2011).

36. C. Y. Ewald, The Matrisome during Aging and Longevity: A Systems-Level Approach toward Defining Matreotypes Promoting Healthy Aging. Gerontology 66, 266–274 (2020).

37. A. C. Teuscher, E. Jongsma, M. N. Davis, C. Statzer, J. M. Gebauer, A. Naba, C. Y. Ewald, The in-silico characterization of the Caenorhabditis elegans matrisome and proposal of a novel collagen classification. Matrix Biol Plus 1, 100001 (2019).

38. A. Sainio, H. Jarvelainen, Extracellular matrix-cell interactions: Focus on therapeutic applications. Cell Signal 66, 109487 (2020).

39. I. Boraschi-Diaz, J. Wang, J. S. Mort, S. V. Komarova, Collagen Type I as a Ligand for Receptor-Mediated Signaling. Front Phys-Lausanne 5, (2017).

40. C. Kumsta, T. T. Ching, M. Nishimura, A. E. Davis, S. Gelino, H. H. Catan, X. Yu, C. C. Chu, B. Ong, S. H. Panowski, N. Baird, R. Bodmer, A. L. Hsu, M. Hansen, Integrin-linked kinase modulates longevity and thermotolerance in C. elegans through neuronal control of HSF-1. Aging Cell 13, 419–430 (2014).

41. D. Kim, B. Langmead, S. L. Salzberg, HISAT: a fast spliced aligner with low memory requirements. Nature methods 12, 357–360 (2015).

42. C. Trapnell, A. Roberts, L. Goff, G. Pertea, D. Kim, D. R. Kelley, H. Pimentel, S. L. Salzberg, J. L. Rinn, L. Pachter, Differential gene and transcript expression analysis of RNA-seq experiments with TopHat and Cufflinks. Nature protocols 7, 562–578 (2012).

